# Multidimensional characterization of the physiological and behavioral effects of TCB-2 in mice

**DOI:** 10.64898/2026.08.01.742216

**Authors:** Mizuki Yamamoto, Hiroto Inoue, Kazuko Hayashi, Illia Aota, Jumpei Matsumoto, Kota Yamada, Koji Toda

## Abstract

**Background and Purpose:** Serotonergic psychedelics affect behavior and physiology, but the relationships among these effects remain poorly understood. In rodents, the head-twitch response is used as a measure of psychedelic-like activity, yet it does not capture changes in physiological state or the performance of learned behaviors. Here, we investigated the acute effects of the 5-HT_2A_ receptor agonist TCB-2 across several behavioral and physiological measures and examined how these effects were modified by pretreatment with the 5-HT_2A_ receptor antagonist volinanserin.

**Experimental Approach:** Mice were tested in head-fixed and freely moving conditions. During a learned auditory trace-conditioning task, we measured licking, pupil area, eye position, and blinking. We measured locomotor activity in an open field and quantified head-twitch responses using a DeepLabCut-based method. To examine the contribution of 5-HT_2A_ receptors, mice were pretreated with the 5-HT_2A_ receptor antagonist volinanserin.

**Key Results:** TCB-2 caused pupil constriction without detectable changes in eye position or blinking when administered alone. TCB-2 also reduced licking at the highest dose, but the cue-locked temporal pattern of licking remained evident. In freely moving mice, TCB-2 reduced locomotor activity and produced a dose-dependent increase in head-twitch responses. Volinanserin partially attenuated TCB-2-induced pupil constriction and reduced head-twitch responses under some conditions, but it did not consistently prevent the other effects of TCB-2.

**Conclusions and Implications:** TCB-2 produced distinct effects across physiological and behavioral measures rather than a uniform disruption of behavioral function. Pronounced pupil constriction and head-twitch responses occurred without detectable changes in eye position or blinking, while the temporal organization of conditioned licking was retained despite a reduction in its magnitude. The incomplete and variable effects of volinanserin preclude definitive conclusions about the receptor mechanisms underlying each response. Combining automated head-twitch detection with physiological and task-related measurements provides a broader framework for comparing the pharmacological profiles of serotonergic compounds.

**What is already known:**

- Classical psychedelics produce characteristic effects primarily through serotonin 5-HT_2A_ receptor activation.
- Head-twitch responses capture only one dimension of psychedelic-like drug action.

**What this study adds:**

- TCB-2 reduced locomotion and licking while preserving the cue-locked pattern of conditioned licking.
- Pupil constriction occurred without detectable changes in eye position or blinking.

**Clinical significance:**

- Multidimensional phenotyping can distinguish the physiological and behavioral profiles of serotonergic compounds.
- Complementary measures may improve preclinical evaluation of emerging serotonergic therapeutics.

## 1. Introduction

Major depressive disorder (MDD) is a major global health problem that causes substantial personal and socioeconomic burdens (GBD 2019 Mental Disorders Collaborators, 2022). Available antidepressants are effective for many patients, but some patients do not achieve remission, and clinical improvement may take several weeks. New treatments that act rapidly and produce lasting benefits are therefore needed (Cipriani et al., 2018; Yamamoto et al., 2026a). Classical psychedelics, particularly psilocybin, are being investigated as potential treatments for psychiatric disorders, including treatment-resistant depression. Clinical studies suggest that psilocybin administered with psychological support can rapidly reduce depressive symptoms and that these improvements may persist after the acute drug effects have subsided (Carhart-Harris et al., 2016, 2018; Davis et al., 2021; Goodwin et al., 2022). The serotonin 5-HT_2A_ receptor is a G protein-coupled receptor that is widely expressed in cortical regions involved in perception, cognition, and emotion. It is a principal pharmacological target of classical psychedelics and contributes to their acute subjective effects, with a possible role in their therapeutic effects (Sharp & Barnes, 2020; Vollenweider & Preller, 2020). However, it remains unclear how the pharmacological actions of serotonergic psychedelics at this receptor relate to their distinct neural, physiological, and behavioral effects.

Compounds that act at 5-HT_2A_ receptors can alter several aspects of neural function, including cortical excitability, large-scale network activity, sensory processing, and synaptic plasticity (de Araujo et al., 2012; Kometer et al., 2013; Shao et al., 2021, 2025). These neural effects may contribute both to the acute subjective experiences produced by psychedelics and to changes that persist after the acute drug effects have subsided (Carhart-Harris & Friston, 2019). Psychedelics may alter perception and cognition by changing how the brain processes and integrates external sensory information with internally generated representations (Alamia et al., 2020; Duerler et al., 2022; Roseman et al., 2016). However, it remains unclear how the acute pharmacological effects of serotonergic psychedelics relate to changes in physiological state, responses to sensory cues, and engagement in ongoing behavior. Measuring several behavioral and physiological responses, as well as their sensitivity to receptor antagonism, may provide a more complete description of the pharmacological profile of each compound.

Rodent models are widely used to study the behavioral and pharmacological effects of psychedelic compounds. The head-twitch response is one of the most established behavioral measures associated with 5-HT_2A_ receptor activation (Halberstadt & Geyer, 2018). Classical psychedelics reliably induce head-twitch responses in rodents, and these responses are reduced by 5-HT_2A_ receptor antagonists or by genetic deletion of the receptor (González-Maeso et al., 2007; Jaster et al., 2022; Takaba et al., 2024). The potency of serotonergic psychedelics in producing head-twitch responses in mice also correlates with their behavioral and subjective potency in other species, supporting the translational value of this assay (Halberstadt et al., 2020). However, the head-twitch response is a single unconditioned behavior. It does not show whether a compound also alters physiological state, spontaneous activity, sensory processing, or the performance of a learned task (Canal & Morgan, 2012). Additional measures are therefore needed to characterize the broader effects of serotonergic compounds.

TCB-2 is a conformationally restricted phenethylamine with high affinity and agonist activity at the 5-HT_2A_ receptor. It can activate different receptor-mediated signaling pathways to different degrees, making it a useful pharmacological tool for studying 5-HT_2A_ receptor function (McLean et al., 2006). In C57BL/6J mice, systemic administration of TCB-2 produces dose-dependent head-twitch responses, hypothermia, reduced food intake, and increased corticosterone concentrations. Several of these effects are reduced by 5-HT_2A_ receptor antagonism (Fox et al., 2010). However, studies have reported different effects of TCB-2 on locomotor activity. Fox et al. (2010) found no significant change in open-field activity, whereas Halberstadt et al. (2013) observed locomotor stimulation that was absent in 5-HT_2A_ receptor-knockout mice and blocked by M100907. These differences may reflect variation in dose, habituation, testing conditions, session duration, or the time period used for analysis. Most previous studies of TCB-2 have focused on unconditioned behavioral and physiological responses. It remains unclear whether acute TCB-2 administration disrupts the performance of a previously learned, sensory-guided behavior and whether volinanserin modifies these effects.

Answering these questions requires simultaneous measurement of learned behavior and physiological state. Pupil dynamics provide a noninvasive measure related to arousal, neuromodulatory activity, sensory processing, and task engagement (Joshi & Gold, 2020; Reimer et al., 2016). However, pupil measurements alone cannot show whether a drug-induced change is limited to pupil control or occurs alongside changes in eye position, blinking, or engagement in the task. Monitoring eye position and blinking provides additional information needed to interpret changes in pupil dynamics and task performance. Measuring these variables while recording licking during a sensory-guided task can help determine whether a drug changes physiological state without causing a broad disruption of motor output, eye stability, eyelid activity, or task engagement. This approach is especially relevant to serotonergic compounds because their effects on perception and behavioral state may differ across physiological and behavioral measures.

In the present study, we characterized the acute behavioral and physiological effects of TCB-2 in mice and examined how volinanserin pretreatment modified these effects. We used freely moving and head-fixed paradigms to measure several aspects of the drug response. In freely moving mice, we measured open-field activity, fecal and urinary output, and head-twitch responses. Head-twitch responses were quantified using an automated video-based computer-vision method. In head-fixed mice, we measured licking, pupil dynamics, eye position, and blinking during a previously learned auditory trace-conditioning task. This combination of measurements allowed us to compare the effects of TCB-2 on spontaneous activity, learned task performance, pupil state, eye position, blinking, and head-twitch responses. We specifically asked whether the marked behavioral and physiological effects of TCB-2 were accompanied by a broad disruption of eye stability, blinking, task engagement, or the temporal organization of learned behavior, and whether volinanserin altered each response.

## 2. Methods

### 2.1. Animals and ethical approval

A total of fifty adult male C57BL/6J mice were used in this study. Mice were experimentally naive at the beginning of the study. Animals were maintained on a reversed 12-h light–dark cycle (lights on at 20:00) in a temperature-controlled room (24 ± 2°C). Experiments were conducted during the dark phase, and individual mice were tested at approximately the same time of day whenever possible. Before surgery, mice were housed in groups of two to four per cage. Food and water were available ad libitum except during the water-restriction procedure described below. Procedures were approved by the Animal Care and Use Committee of Keio University. Animal health and body weight were monitored throughout the study.

### 2.2. Experimental design

Separate cohorts were used for the head-fixed auditory trace-conditioning task, open-field task, and head-twitch response test. The experimental unit was an individual mouse. Treatment orders were randomized within each experiment. Group sizes were determined on the basis of previous studies and established laboratory protocols. No formal sample-size calculation was performed. For experiments with repeated drug administration, each mouse received all treatments assigned to its cohort in a within-subject design. Treatment order was randomized, and successive test sessions were separated by at least 1 day. The exact number of animals and independent observations included in each analysis is reported in the corresponding figure legend.

### 2.3. Surgery

Mice assigned to the head-fixed experiments underwent surgery for implantation of a head plate. Anaesthesia was induced and maintained with 1.0–2.5% isoflurane in room air. Mice were placed in a stereotaxic frame (model 942WOAE; David Kopf Instruments, Tujunga, CA, USA). Following shaving, the skull was exposed and a head plate (H.E. Parmer Company, Nashville, TN, USA) was secured to the skull using dental cement (product no. 56849; 3M, St Paul, MN, USA). After surgery, mice were housed individually and allowed to recover for at least 7 days before water restriction and behavioral training began. Postoperative health, body weight, and the condition of the surgical site were monitored daily.

### 2.4. Drugs and administration

TCB-2 was purchased from Tocris Bioscience (Bristol, UK) and dissolved in physiological saline. TCB-2 was administered intraperitoneally at 0.3, 1.0, or 3.0 mg/kg in an injection volume of 10 mL/kg. Control animals received an equivalent volume of saline. The dose range was selected on the basis of previous studies examining the behavioral effects of TCB-2 (Fox et al., 2010; Rahbarnia et al., 2025). Behavioral testing began 5 min after TCB-2 or saline administration. Volinanserin (MDL 100,907) was purchased from Selleck Chemicals (Houston, TX, USA) and dissolved in physiological saline containing 1% dimethyl sulfoxide (DMSO). Volinanserin was administered intraperitoneally at 1.0 mg/kg in an injection volume of 10 mL/kg. The dose was selected on the basis of previous pharmacological studies of 5-HT_2A_ receptor antagonism (Takaba et al., 2024). Vehicle controls received physiological saline containing 1% DMSO. In the antagonist experiments, vehicle or volinanserin was administered 30 min before saline or TCB-2. TCB-2 and volinanserin solutions were prepared in advance, stored at −80°C, and thawed immediately before use.

### 2.5. Auditory trace-conditioning task

#### 2.5.1. Animals, apparatus, and training

Sixteen mice were used in the head-fixed experiments. Eight mice were assigned to the TCB-2-only cohort (age range, 2.8–4.3 months; mean ± SD, 3.5 ± 0.5 months), and eight were assigned to the volinanserin-pretreatment cohort (age range, 3.0–6.5 months; mean ± SD, 3.75 ± 1.6 months). After postoperative recovery, mice were water-restricted for 2 days before behavioral training began. Body weight was measured daily and maintained above 85% of each animal’s free-drinking weight. A 10% sucrose solution was provided during training and testing, and supplementary water was provided after each session when required. Food remained available ad libitum.

Behavioral testing was performed in a square chamber. Mice were head-fixed on a custom-designed, 3D-printed platform equipped with a camera for eye recording (Fig. 1A). The apparatus and behavioral procedures were adapted from previously described head-fixed conditioning paradigms (Toda et al., 2017; Yamada & Toda, 2022; Yamada et al., 2024; Yamamoto et al., 2022; Inoue et al., 2026). A stainless-steel drinking spout was positioned directly in front of the mouth. Licks were detected using a contact lickometer connected between the drinking spout and a metal plate beneath the platform. Reward delivery and lick recording were controlled using custom-written Python scripts (Python 3.7.7) and a custom relay circuit connected to solenoid valves.

**Figure 1.**
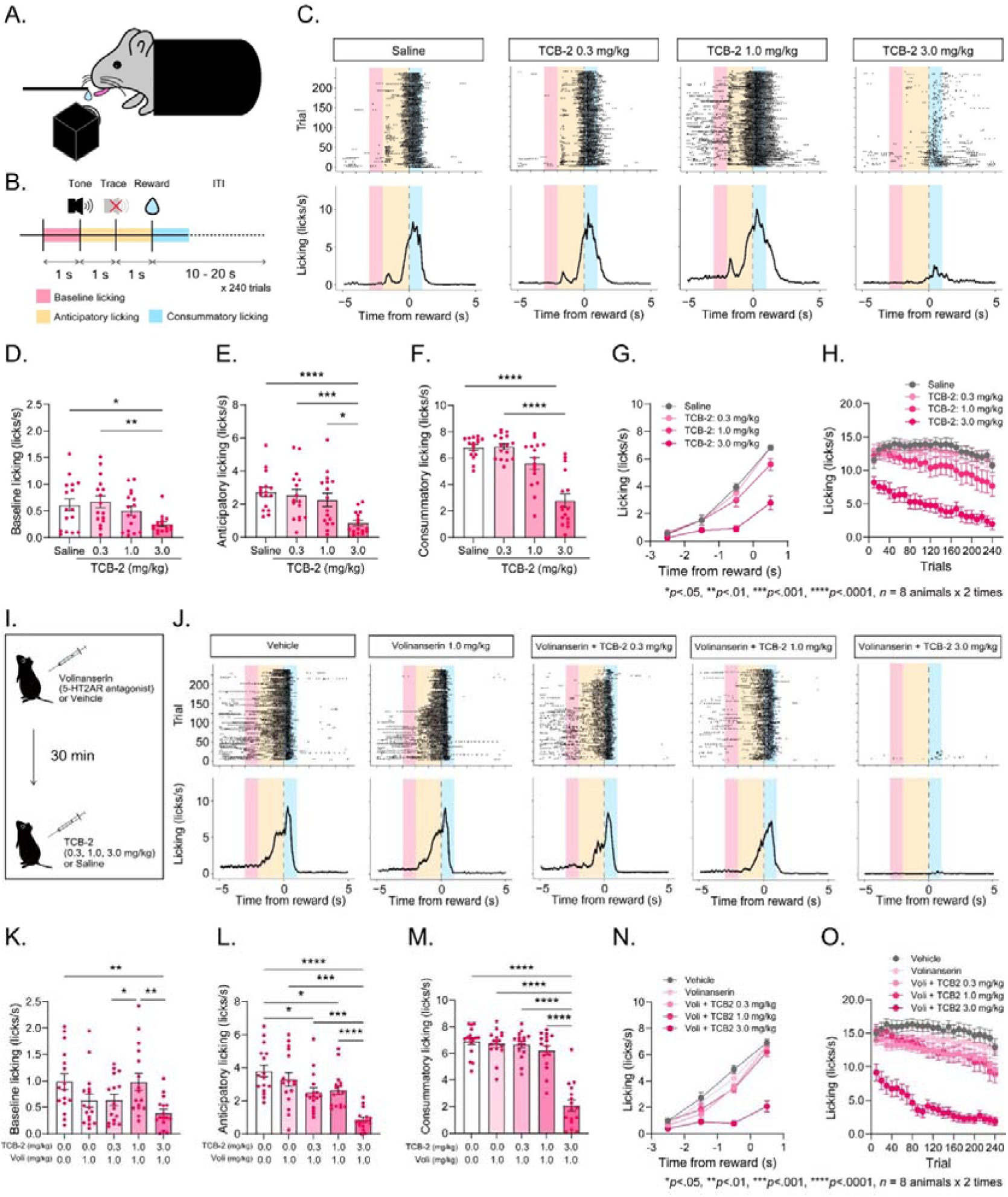
Effects of TCB-2 on licking responses during auditory trace conditioning with and without volinanserin pretreatment. (A) Schematic side view of a head-fixed mouse during the task. (B) Timeline of the auditory trace-conditioning task. (C) Representative licking responses from a single mouse following administration of saline or TCB-2 at 0.3, 1.0, or 3.0 mg/kg (from left to right). The upper and lower panels show lick rasters and lick-density plots, respectively. (D) Effects of TCB-2 on baseline licking. (E) Effects of TCB-2 on anticipatory licking. (F) Effects of TCB-2 on consummatory licking. (G) Effects of TCB-2 on licking across the task timeline. (H) Effects of TCB-2 on licking over the course of successive trials. (I) Schematic timeline of volinanserin pretreatment and subsequent TCB-2 administration. (J) Representative licking responses from a single mouse following administration of vehicle, volinanserin alone (1.0 mg/kg), or volinanserin (1.0 mg/kg) followed by TCB-2 at 0.3, 1.0, or 3.0 mg/kg (from left to right). The upper and lower panels show lick rasters and lick-density plots, respectively. (K) Effects of TCB-2 following volinanserin pretreatment on baseline licking. (L) Effects of TCB-2 following volinanserin pretreatment on anticipatory licking. (M) Effects of TCB-2 following volinanserin pretreatment on consummatory licking. (N) Effects of TCB-2 following volinanserin pretreatment on licking across the task timeline. (O) Effects of TCB-2 following volinanserin pretreatment on licking over the course of successive trials. Data were obtained from eight mice, each tested twice under each condition. Error bars represent the SEM. \**p* < 0.05, \*\**p* < 0.01, \*\*\**p* < 0.001, and \*\*\*\**p* < 0.0001.

Each trial consisted of a 1-s auditory cue, followed by a 1-s trace interval and delivery of a 10% sucrose reward (Fig. 1B). The auditory cue was presented at 6 kHz and 80 dB. The volume and duration of sucrose delivery were approximately 4.2 μL and 10 ms, respectively. Intertrial intervals varied pseudorandomly between 10 and 20 s, and each training or testing session comprised 240 trials. Baseline licking was defined as the number of licks during the 1 s preceding tone onset. Anticipatory licking was defined as the number of licks during the 1-s tone and the subsequent 1-s trace interval. Consummatory licking was defined as the number of licks during the first second after reward delivery. Each mouse completed two test sessions under each treatment condition. Treatment order was randomized, and successive test sessions were separated by at least 1 day.

#### 2.5.2. Pupil, eye-position, and blink measurements

The eye was recorded while mice performed the auditory trace-conditioning task. An infrared camera was positioned 45 mm from the top of the head at an angle of 50° relative to the animal’s midline. Infrared illumination was used to visualize the pupil, and visible illuminance inside the chamber was maintained at 95 lux.

Pupil and eyelid landmarks were tracked using DeepLabCut (Mathis et al., 2018), following the general approach described by Yamada and Toda (2022). The trained network identified the upper, lower, left, and right eyelid margins and 8 landmarks along the pupil boundary. An ellipse was fitted to the tracked pupil-boundary landmarks in each frame. Pupil area was calculated from the fitted major and minor axes. Pixel measurements were converted to physical units. The center of the fitted ellipse was used to estimate horizontal and vertical eye position. Changes in eye position were calculated. Overall pupil area was calculated as mean pupil area across the complete session. For task-aligned analyses, pupil measurements were aligned to tone onset and averaged across valid trials.

### 2.6. Open-field task

Eighteen mice were used in the open-field experiments. Ten mice were assigned to the TCB-2-only cohort (age range, 2.6–5.7 months; mean ± SD, 3.6 ± 0.8 months), and eight were assigned to the volinanserin-pretreatment cohort (age range, 4.3–6.5 months; mean ± SD, 5.9 ± 1.0 months). The procedure was adapted from previously described open-field protocols (Kaneko et al., 2022, 2026; Tamura et al., 2024; Ujihara et al., 2024; Yamamoto et al., 2026b; Inoue et al., 2026).

The open-field arena was a custom-made white polyvinyl chloride box measuring 50 × 50 × 50 cm. A camera (HD Pro Webcam C920r; Logicool, Tokyo, Japan) was positioned 106 cm above the floor of the arena, and videos were acquired at 30 frames/s using a Windows computer. White noise was presented continuously at 75 dB to mask external sounds.

Before drug testing, mice underwent three daily 60-min habituation sessions. At the beginning of each habituation session, mice received an intraperitoneal saline injection and were immediately placed in the arena. During test sessions, mice received saline or TCB-2 at 0.3, 1.0, or 3.0 mg/kg and were placed in the arena 5 min later. Each session lasted 60 min. Treatment order was randomized within animals, and test sessions were separated by at least 3 days. For the antagonist experiment, mice received vehicle or volinanserin 30 min before saline or TCB-2, and open-field testing began 5 min after the second injection. After each session, the arena was cleaned with 70% ethanol and allowed to dry for at least 15 min. The investigator conducting the subsequent session was not blinded to treatment conditions.

Mouse position was tracked using Bonsai (Lopes et al., 2015). Video frames were converted to greyscale, smoothed, and intensity-inverted. The animal was segmented using a predefined contrast threshold, and the centroid coordinates were calculated for each frame. Locomotor activity was expressed as distance travelled per unit time and was calculated from changes in centroid position after conversion from pixels to centimetres. The center region was defined as the central 25 × 25 cm area of the open-field arena. Center time was calculated as the cumulative duration for which the animal’s centroid remained within this region. Locomotor activity and center time were calculated for the complete 60-min session and in 5-min time bins. Immediately after each session, fecal boli were collected and weighed. Investigators performing these measurements were not blinded to treatment conditions.

### 2.7. Head-twitch response test

#### 2.7.1. Experimental procedure

Sixteen mice were allocated to the head-twitch response experiment. Eight mice were assigned to the TCB-2-only cohort (age range, 3.2–3.9 months; mean ± SD, 3.8 ± 0.8 months), and eight mice were assigned to the volinanserin-pretreatment cohort (age range, 3.6–6.6 months; mean ± SD, 4.67 ± 0.9 months). Testing was performed in a custom-made cylindrical white polyvinyl chloride chamber measuring 20 cm in diameter and 30 cm in height. A GoPro HERO13 Black camera (GoPro Inc., San Mateo, CA, USA) was positioned 30 cm above the floor of the chamber. Mice in the TCB-2-only cohort were allowed to habituate to the chamber for 15 min and then received saline or TCB-2 at 0.3, 1.0, or 3.0 mg/kg. Mice in the antagonist cohort received vehicle or volinanserin 30 min before saline or TCB-2. Recording began 5 min after the second injection and continued for 60 min at 120 frames/s. Each mouse received randomized order treatment, with an interval of at least 1 day between sessions.

#### 2.7.2. Automated detection of head-twitch responses

Head-twitch responses were quantified using an automated detection system that combined DeepLabCut-based pose estimation with a support vector machine (SVM) classifier. At least 1 h before behavioral testing, the right and left ears of each mouse were marked with pink and blue paint, respectively. Videos were recorded at 120 frames/s with sufficient image quality for markerless pose estimation using DeepLabCut (Mathis et al., 2018). Data used to develop the detection system were obtained from four recording sessions selected across the saline and TCB-2 treatment conditions (0.3, 1.0, and 3.0 mg/kg). Head-twitch events in these recordings were manually annotated using BORIS version 9.7.15 (Friard & Gamba, 2016). DeepLabCut was used to track the left ear, right ear, nose, body center, and tail base. The manually annotated event timestamps were then synchronized with the tracked coordinate data to generate labeled examples for classifier development.

Nine features were calculated from the tracked body-part coordinates. Eight features represented high-frequency power in the 20–60 Hz range. These features were obtained by applying a wavelet transform to the x- and y-coordinates of the left ear, right ear, nose, and body center in an animal-centered coordinate system. The ninth feature was the speed of the body center. The high-frequency power derived from the ear and nose coordinates was summed, and time points at which this total power reached a peak exceeding 2 standard deviations above its mean were identified as candidate head-twitch events. Each candidate event was assigned a label based on the manually annotated head-twitch events. The nine DeepLabCut-derived features and the corresponding labels were then used to train an SVM classifier in MATLAB version 24.2.0 (MathWorks, Natick, MA, USA). Classifier performance was evaluated using held-out test segments that were not used for classifier training or parameter selection. Automated detections were compared with manually annotated head-twitch events using a temporal matching tolerance of 5 seconds. Recall was calculated as TP/(TP + FN), precision as TP/(TP + FP), and the F1 score as 2 × precision × recall/(precision + recall), where TP, FP, and FN represent true-positive, false-positive, and false-negative detections, respectively. To assess inter-rater agreement in manual annotation, 10-min recordings obtained from the same eight mice under each of four treatment conditions were independently evaluated by two observers who were blinded to treatment. Each recording was divided into consecutive, non-overlapping 5-s windows. For each observer, every window was assigned a binary score indicating whether at least one head-twitch event was present (1) or absent (0), regardless of the number of events within the window. Binary scores from all recordings were pooled, and Cohen’s κ was calculated to quantify the overall agreement between the two observers.

### 2.8. Data and statistical analysis

Data were analysed using RStudio version 2022.02.0 (RStudio PBC, Boston, MA, USA), GraphPad Prism version 10.4.0 (GraphPad Software, Boston, MA, USA), and MATLAB version 24.2.0 (MathWorks). The data and statistical analysis complied with the recommendations on experimental design and analysis in pharmacology described in the British Journal of Pharmacology guidance. The individual mouse was the experimental unit. Repeated measurements or duplicate sessions obtained from the same mouse were not treated as independent biological replicates. Group sizes reported in the figure legends refer to animals. Data are presented as individual observations with the mean ± SEM unless otherwise stated. Statistical analyses were performed only for groups containing at least five independent animals. No data were removed as statistical outliers. Dose-dependent effects measured repeatedly in the same animals were analysed using repeated-measures one-way ANOVA. When the assumption of sphericity was violated, the Greenhouse–Geisser correction was applied, as indicated by non-integer degrees of freedom. Post hoc comparisons were performed using Tukey’s test. Post hoc testing was conducted only when the overall ANOVA was significant and the assumptions required for the analysis were satisfied. Cohen’s κ was calculated to assess agreement between manual observers. Classifier performance was summarized using recall, precision, and F1 score. These performance measures were calculated from held-out observations and were not subjected to inferential statistical testing. A two-sided P value below 0.05 was considered statistically significant. Exact *p* values are reported wherever possible; values below 0.0001 are reported as *p* < 0.0001.

## 3. Results

### 3.1. Effects of TCB-2 on licking during auditory trace conditioning with and without volinanserin pretreatment

To investigate the effects of TCB-2 on the performance of a learned, sensory-guided behavior, we tested head-fixed mice in an auditory trace-conditioning task (Fig. 1A, B). Following training, all mice successfully acquired the task. Administration of TCB-2 reduced licking at the highest dose tested, as illustrated by the representative lick rasters and lick-density plots (Fig. 1C). This reduction was not restricted to a particular task epoch: separate analyses showed reductions in baseline licking (Fig. 1D; *F*(2.499, 37.48) = 5.455, *p* = 0.0051, repeated-measures one-way ANOVA), anticipatory licking (Fig. 1E; *F*(2.050, 30.75) = 14.17, *p* < 0.0001, repeated-measures one-way ANOVA), and consummatory licking (Fig. 1F; *F*(2.033, 30.50) = 13.60, *p* < 0.0001, repeated-measures one-way ANOVA). Analysis across the task timeline confirmed a broadly distributed reduction in licking rather than an effect confined to a single behavioral epoch (Fig. 1G). Analysis across successive trials showed that the reduction developed over the course of the test session (Fig. 1H).

To evaluate the effect of volinanserin on the TCB-2-induced changes in licking, mice were pretreated with volinanserin 30 min before TCB-2 administration (Fig. 1I). Representative lick rasters and lick-density plots for each treatment condition are shown in Fig. 1J. Volinanserin pretreatment did not prevent the reduction in licking produced by the highest dose of TCB-2. Following pretreatment, reductions remained evident in baseline licking at the highest TCB-2 dose (Fig. 1K; *F*(2.352, 30.58) = 5.510, *p* = 0.0060, repeated-measures one-way ANOVA), in anticipatory licking across the TCB-2-treated conditions (Fig. 1L; *F*(3.077, 46.16) = 21.86, *p* < 0.0001, repeated-measures one-way ANOVA), and in consummatory licking at the highest dose (Fig. 1M; *F*(2.868, 43.02) = 5.455, *p* < 0.0001, repeated-measures one-way ANOVA). Analyses across the task timeline and successive trials further showed that the suppression of licking remained evident following volinanserin pretreatment (Fig. 1N, O).

### 3.2. Effects of TCB-2 on pupil dynamics, eye position, and blinking during auditory trace conditioning with and without volinanserin pretreatment

While mice performed the auditory trace-conditioning task, videos of the eye were recorded under head-fixed conditions. Using DeepLabCut-based landmark tracking (Mathis et al., 2018) and ellipse fitting (Yamada & Toda, 2022), we quantified pupil area and eye position (Fig. 2A, B). TCB-2 decreased overall pupil area (Fig. 2C; *F*(2.241, 33.62) = 18.77, *p* < 0.0001, repeated-measures one-way ANOVA). Analysis across the test session showed that the reduction in pupil area remained evident over time (Fig. 2D). Analysis of pupil dynamics aligned to tone onset showed that pupil area gradually increased between tone presentation and reward delivery (Fig. 2E). This task-related dilation remained evident at 0.3 and 1.0 mg/kg but was attenuated at 3.0 mg/kg following normalization of pupil area (Fig. 2E, F).

**Figure 2.**
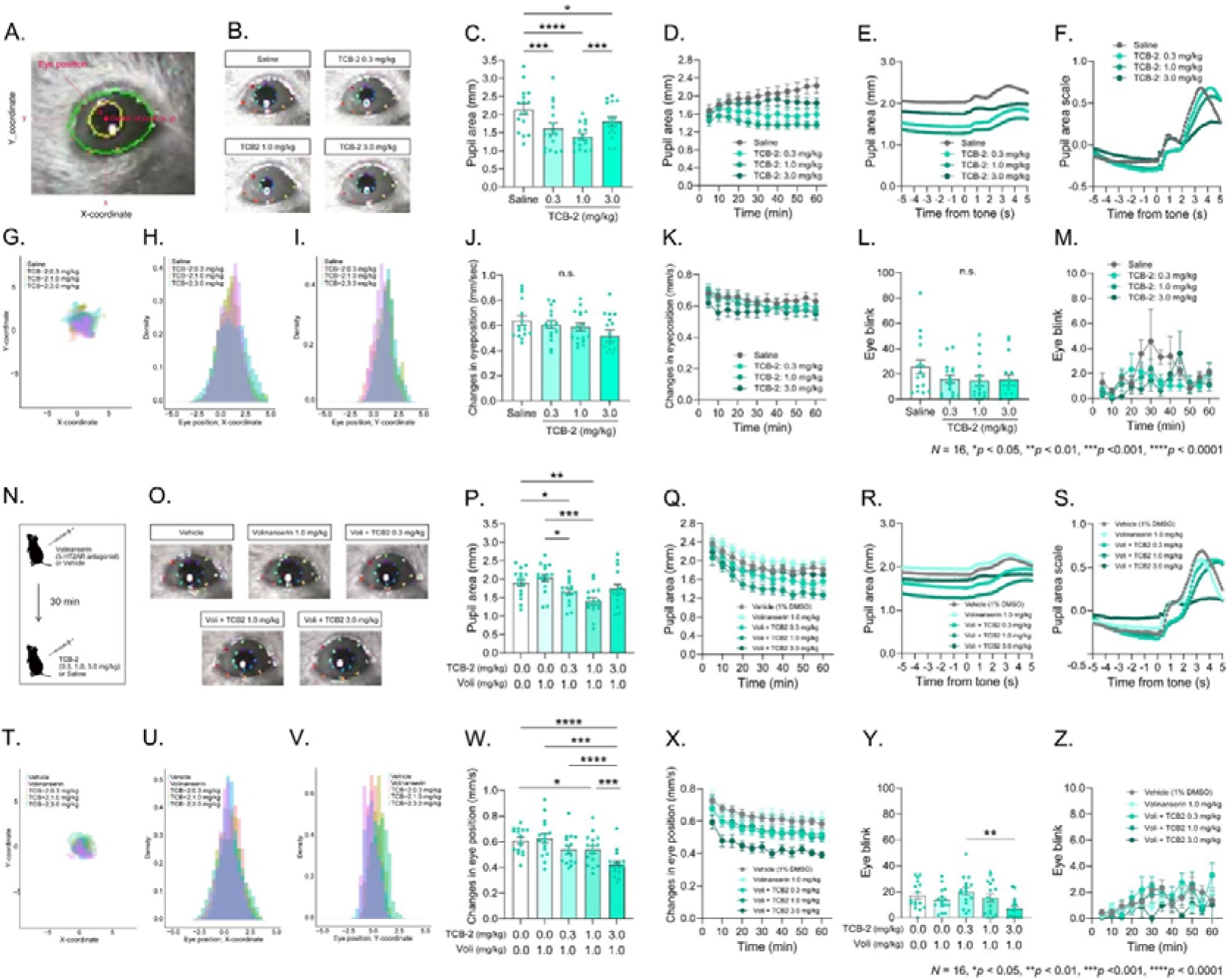
Effects of TCB-2 on pupil dynamics, eye position, and blinking during auditory trace conditioning with and without volinanserin pretreatment. (A) Schematic illustration of pupil-size and eye-position measurements. (B) Representative pupil-size recordings following intraperitoneal administration of saline or TCB-2 at 0.3, 1.0, or 3.0 mg/kg. (C) Effects of TCB-2 on overall pupil size. (D) Average time course of pupil size following TCB-2 administration. (E) Average pupil dynamics aligned to tone onset. (F) Normalized pupil dynamics following TCB-2 administration. (G) Representative eye positions following saline or TCB-2 administration. (H) Distribution of the horizontal coordinates of eye position. (I) Distribution of the vertical coordinates of eye position. (J) Effects of TCB-2 on eye position. (K) Effects of TCB-2 on changes in eye position. (L) Effects of TCB-2 on blinking. (M) Time course of blinking following TCB-2 administration. (N) Schematic timeline of volinanserin pretreatment and subsequent TCB-2 administration. (O) Representative pupil-size recordings following administration of vehicle, volinanserin alone (1.0 mg/kg), or volinanserin (1.0 mg/kg) followed by TCB-2 at 0.3, 1.0, or 3.0 mg/kg. (P) Effects of TCB-2 following volinanserin pretreatment on overall pupil size. (Q) Average time course of pupil size following TCB-2 administration with volinanserin pretreatment. (R) Average pupil dynamics aligned to tone onset following TCB-2 administration with volinanserin pretreatment. (S) Normalized pupil dynamics following TCB-2 administration with volinanserin pretreatment. (T) Representative eye positions following administration of vehicle, volinanserin alone, or TCB-2 with volinanserin pretreatment. (U) Distribution of the horizontal coordinates of eye position following volinanserin pretreatment. (V) Distribution of the vertical coordinates of eye position following volinanserin pretreatment. (W) Effects of TCB-2 following volinanserin pretreatment on eye position. (X) Effects of TCB-2 following volinanserin pretreatment on changes in eye position. (Y) Effects of TCB-2 following volinanserin pretreatment on blinking. (Z) Time course of blinking following TCB-2 administration with volinanserin pretreatment. Data were obtained from eight mice, each tested twice under each condition. Error bars represent the SEM. n.s., not significant; \**p* < 0.05, \*\**p* < 0.01, \*\*\**p* < 0.001, and \*\*\*\**p* < 0.0001.

The spatial distributions of horizontal and vertical eye position were broadly comparable across treatment conditions (Fig. 2G–I), and TCB-2 did not significantly alter the overall magnitude or time course of changes in eye position (Fig. 2J, K; *F*(1.752, 26.27) = 2.707, *p* = 0.0914). TCB-2 also did not significantly affect the total number of blinks (Fig. 2L; *F*(2.438, 36.57) = 1.688, *p* = 0.1978, repeated-measures one-way ANOVA) or their temporal distribution across the session (Fig. 2M).

Following volinanserin pretreatment (Fig. 2N), TCB-2 continued to reduce overall pupil area (Fig. 2O, P), and this reduction remained evident across the test session (Fig. 2Q). Tone-related pupil dilation remained evident at the lower TCB-2 doses but was attenuated at 3.0 mg/kg (Fig. 2R, S). The spatial distributions of horizontal and vertical eye position remained broadly overlapping across treatment conditions (Fig. 2T–V). However, the highest dose of TCB-2 reduced the magnitude of changes in eye position following volinanserin pretreatment (Fig. 2W, X; *F*(2.257, 33.86) = 18.70, *p* < 0.0001). The total number of blinks also differed among the pretreatment conditions, with fewer blinks observed following the highest dose of TCB-2 (Fig. 2Y; *F*(2.580, 38.70) = 3.645, *p* = 0.0258, repeated-measures one-way ANOVA), although no clear treatment-specific temporal pattern was observed across the session (Fig. 2Z).

### 3.3. Open-field task

To examine the effects of TCB-2 on spontaneous locomotor activity, mice were tested in an open-field arena beginning 5 min after drug administration (Fig. 3A–D). Administration of TCB-2 decreased locomotor activity at the highest dose tested (Fig. 3E; *F*(2.324, 20.92) = 11.58, *p* = 0.0003, repeated-measures one-way ANOVA). Time-course analysis showed that this reduction was most pronounced during the early phase of the session and gradually diminished thereafter (Fig. 3F). In contrast, TCB-2 did not significantly affect the total time spent in the center area (Fig. 3I; *F*(1.909, 17.18) = 0.5335, *p* = 0.5879, repeated-measures one-way ANOVA), although some temporal variation was observed during the session (Fig. 3J).

**Figure 3.**
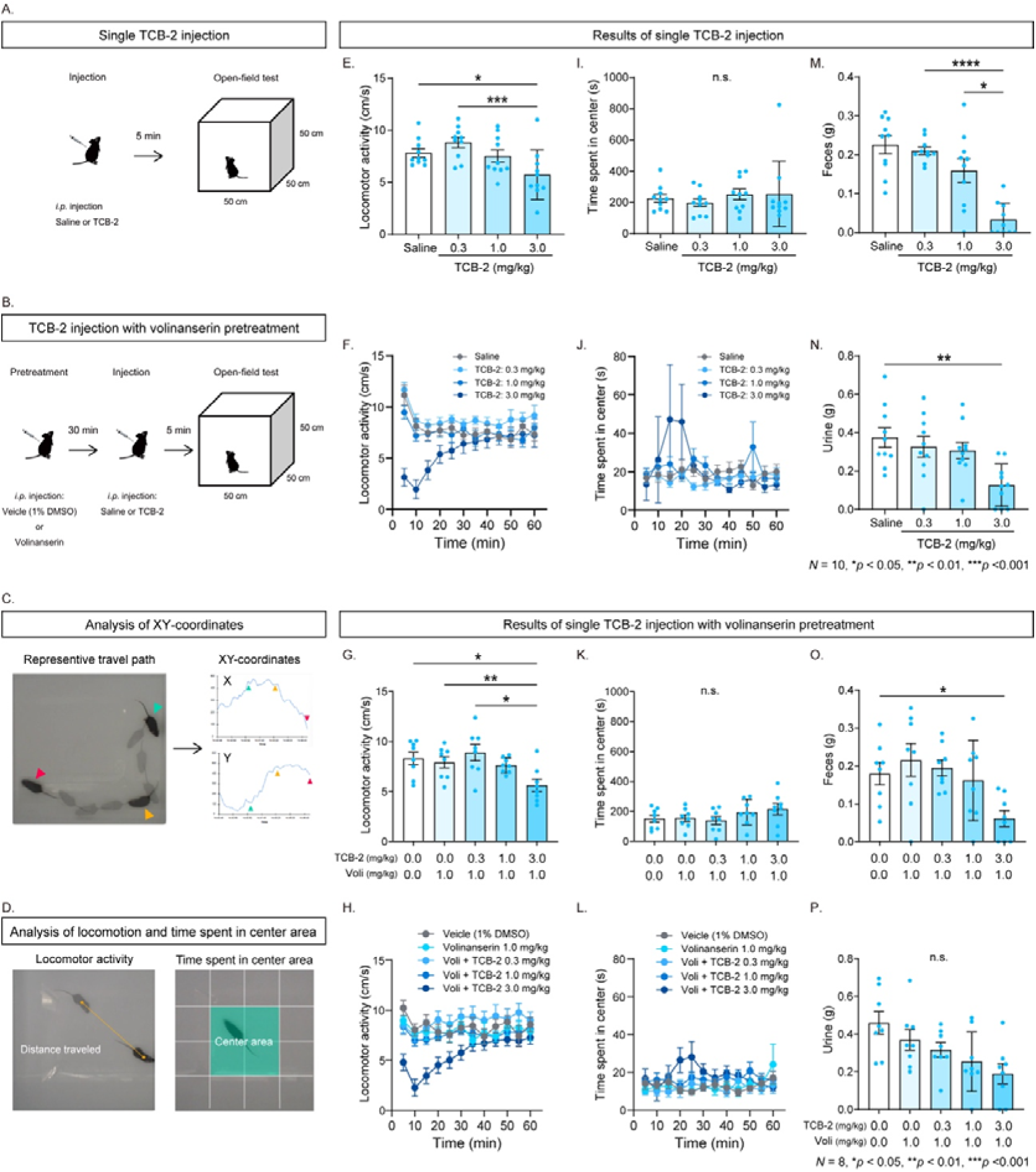
Effects of TCB-2 on open-field activity with and without volinanserin pretreatment. (A) Experimental timeline for the TCB-2-only experiment. Mice received an intraperitoneal injection of saline or TCB-2 and were placed in the open-field arena 5 min later. (B) Experimental timeline for the volinanserin-pretreatment experiment. Mice received an intraperitoneal injection of vehicle or volinanserin (1.0 mg/kg), followed 30 min later by saline or TCB-2. Open-field testing began 5 min after the second injection. (C) Representative travel path and corresponding horizontal and vertical coordinates obtained during the open-field test. (D) Illustration of the measurements used to quantify locomotor activity and time spent in the center area. (E) Mean locomotor activity during the 60-min session following administration of saline or TCB-2 at 0.3, 1.0, or 3.0 mg/kg. (F) Time course of locomotor activity following TCB-2 administration. (G) Mean locomotor activity following administration of vehicle, volinanserin alone, or volinanserin followed by TCB-2 at 0.3, 1.0, or 3.0 mg/kg. (H) Corresponding time course of locomotor activity following volinanserin pretreatment. (I) Total time spent in the center area following TCB-2 administration. (J) Time course of time spent in the center area following TCB-2 administration. (K) Total time spent in the center area following volinanserin pretreatment. (L) Corresponding time course of time spent in the center area following volinanserin pretreatment. (M) Fecal output collected after open-field testing following TCB-2 administration. (N) Urinary output collected after open-field testing following TCB-2 administration. (O) Fecal output collected after open-field testing following volinanserin pretreatment. (P) Urinary output collected after open-field testing following volinanserin pretreatment. Data were obtained from 10 mice in the TCB-2-only experiment and eight mice in the volinanserin-pretreatment experiment. Error bars represent the SEM. Voli, volinanserin. n.s., not significant; \**p* < 0.05, \*\**p* < 0.01, \*\*\**p* < 0.001, and \*\*\*\**p* < 0.0001.

Volinanserin pretreatment did not prevent the reduction in locomotor activity produced by the highest dose of TCB-2 (Fig. 3G; *F*(2.447, 17.13) = 12.66, *p* = 0.0002, repeated-measures one-way ANOVA). Time-course analysis similarly revealed a marked reduction during the early phase of the session, followed by partial recovery (Fig. 3H). Under the volinanserin-pretreatment conditions, there was a significant overall effect of treatment on the total time spent in the center area (Fig. 3K; *F*(3.112, 21.78) = 3.597, *p* = 0.0287, repeated-measures one-way ANOVA). However, post hoc multiple-comparison testing identified no significant differences between individual treatment conditions. The temporal profile of time spent in the center area remained broadly comparable across treatment conditions (Fig. 3L).

To determine whether TCB-2 affected fecal and urinary output during the open-field task, feces and urine were collected and quantified after testing. Administration of the highest dose of TCB-2 decreased both fecal output (Fig. 3M; *F*(1.885, 16.97) = 20.73, *p* < 0.0001, repeated-measures one-way ANOVA) and urinary output (Fig. 3N; *F*(2.756, 24.81) = 6.515, *p* = 0.0026, repeated-measures one-way ANOVA). Following volinanserin pretreatment, the reduction in fecal output at the highest dose of TCB-2 remained evident (Fig. 3O; *F*(2.428, 17.00) = 5.510, *p* = 0.0109, repeated-measures one-way ANOVA). A significant overall effect of treatment on urinary output was also detected (Fig. 3P; *F*(2.816, 19.71) = 3.466, *p* = 0.0382, repeated-measures one-way ANOVA); however, post hoc multiple-comparison testing did not identify significant differences between individual treatment conditions.

### 3.4. Head-twitch response test

To characterize the effects of TCB-2 on head-twitch responses, mice were recorded from above, and head-twitch events were quantified using a custom-developed video-based system that combined DeepLabCut-based pose estimation with an SVM classifier (Fig. 4A, B; see Methods). In held-out test segments, the automated system achieved a recall of 0.972, a precision of 0.882, and an F1 score of 0.925. Manual annotations showed high inter-rater agreement between the two observers (Cohen’s κ = 0.9196).

**Figure 4.**
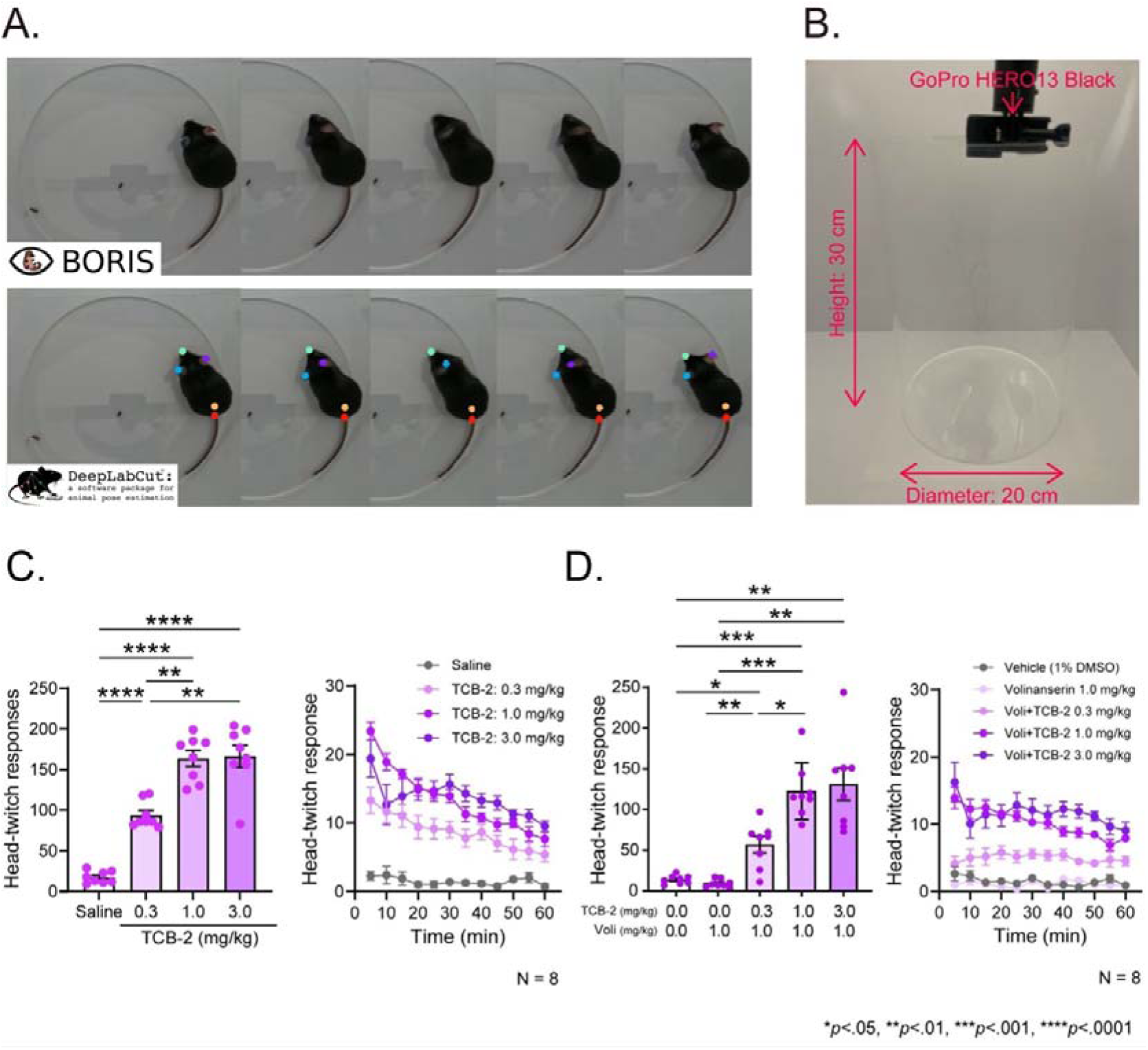
Automated video-based quantification of TCB-2-induced head-twitch responses with and without volinanserin pretreatment. (A) Representative video frames of a head-twitch response scored manually using BORIS (upper panels) and the corresponding anatomical landmarks tracked using DeepLabCut (lower panels). (B) Overhead video-recording setup. Mice were placed individually in a transparent cylindrical arena with a diameter of 20 cm, and behavior was recorded using a GoPro HERO13 Black camera positioned 30 cm above the arena. (C) Effects of TCB-2 on head-twitch responses. The left panel shows the total number of head-twitch responses recorded during the 60-min observation period following administration of saline or TCB-2 at 0.3, 1.0, or 3.0 mg/kg. The right panel shows the corresponding time course of head-twitch responses. (D) Head-twitch responses following volinanserin pretreatment. The left panel shows the total number of responses following administration of vehicle, volinanserin alone (1.0 mg/kg), or volinanserin (1.0 mg/kg) followed by TCB-2 at 0.3, 1.0, or 3.0 mg/kg. The right panel shows the corresponding time course. Data were obtained from eight mice in the TCB-2-only experiment and eight mice in the volinanserin-pretreatment experiment. Error bars represent the SEM. \**p* < 0.05, \*\**p* < 0.01, \*\*\**p* < 0.001, and \*\*\*\**p* < 0.0001.

Intraperitoneal administration of TCB-2 markedly increased the total number of head-twitch responses in a dose-dependent manner (Fig. 4C; *F*(1.974, 13.82) = 78.19, *p* < 0.0001, repeated-measures one-way ANOVA). Time-course analysis showed that the responses emerged immediately after TCB-2 administration and persisted throughout the 60-min observation period (Fig. 4C).

Volinanserin alone did not increase head-twitch responses. Pretreatment with volinanserin attenuated the responses induced by the low dose of TCB-2 (Fig. 4D; *F*(1.470, 10.29) = 30.45, *p* < 0.0001, repeated-measures one-way ANOVA). Nevertheless, TCB-2 continued to produce dose-dependent increases in head-twitch responses following volinanserin pretreatment, and these responses persisted across the observation period (Fig. 4D). Thus, at the dose tested, volinanserin attenuated but did not abolish the head-twitch responses induced by TCB-2.

## 4. Discussion

The present study examined the acute effects of TCB-2 by measuring Pavlovian conditioned behavior, pupil dynamics, eye position, and blinking in head-fixed mice, as well as locomotor activity and head-twitch responses in freely moving mice. TCB-2 reduced pupil area, licking, locomotor activity, and fecal and urinary output, while markedly increasing head-twitch responses. However, these changes did not reflect a uniform disruption across all measures. The cue-locked temporal pattern of licking remained evident despite the reduction in licking, and TCB-2 alone produced no statistically detectable changes in eye-position dynamics or blinking. In a separate experiment involving volinanserin pretreatment, several effects of TCB-2 remained evident, and the highest dose of TCB-2 reduced eye-position changes and blinking. Thus, TCB-2 produced a distinct pattern of effects across behavioral and physiological measures. Because volinanserin did not consistently prevent these effects, the present experiments do not establish the receptor mechanisms underlying each response.

One of the main findings was that TCB-2 produced pronounced pupillary constriction. Pupil dynamics are related to arousal, neuromodulatory activity, sensory processing, and engagement in a task (Joshi & Gold, 2020; Reimer et al., 2016). In head-fixed mice performing a Pavlovian conditioning task, pupil responses also vary with reward prediction and consumption (Yamada & Toda, 2022). In the present study, TCB-2 reduced overall pupil size. However, cue-related pupil dilation remained evident at 0.3 and 1.0 mg/kg and was attenuated only at 3.0 mg/kg. These findings indicate that TCB-2 had different effects on overall pupil size and the pupil response associated with task events. The pupillary constriction observed in mice contrasts with findings in humans, in whom LSD and psilocybin generally produce pupil dilation (Holze et al., 2022). Direct comparison is difficult because pupil responses can differ according to species, compound, dose, illumination, behavioral state, and autonomic control. Even so, this difference shows that serotonergic compounds should be evaluated individually rather than assumed to produce the same pattern of pupil responses.

The pronounced effect of TCB-2 on pupil size, despite the relative stability of eye position and blinking, is particularly relevant to psychedelic research. Classical psychedelics are usually discussed in terms of their effects on cortical activity, perception, and subjective experience, while their effects on eye movements and blinking have received less attention. A recent human eye-tracking study found that psilocybin altered the distribution and timing of gaze fixations during visual exploration, showing that psychedelic effects can be detected through quantitative measurements of eye movements (Muller et al., 2025). In the present study, TCB-2 produced pronounced pupillary constriction during the auditory task and robust head-twitch responses in freely moving mice. However, TCB-2 alone produced no statistically detectable changes in eye-position dynamics or blinking. Direct comparison with human psychedelic states is difficult because of differences in species, compound, dose, visual environment, and behavioral demands. Stable eye position also does not show that sensory processing was unaffected, because changes in perception or sensory processing may occur without an overt change in eye position. Nevertheless, measuring pupil size, eye position, and blinking simultaneously helped determine whether the pupillary constriction was accompanied by a broader change in eye movements or eyelid activity. The stability of eye position makes it less likely that the apparent constriction resulted from gross eye displacement, while the absence of a detectable change in blinking argues against generalized eyelid closure as an explanation.

In the separate experiment involving volinanserin pretreatment, the highest dose of TCB-2 reduced eye-position changes and blinking. These results indicate that pupil size, eye-position dynamics, and blinking do not necessarily change in parallel across treatment conditions. However, the experiments with and without volinanserin pretreatment involved independent cohorts. The difference between the experiments therefore does not show that volinanserin revealed a specific effect or mechanism. In addition, the absence of statistically significant effects on eye position and blinking following TCB-2 alone should not be interpreted as evidence that these measures were unchanged. The precision and statistical power of the present experiments may have been insufficient to detect small effects.

The results of the auditory trace-conditioning task provide another example of how TCB-2 affected different aspects of behavior to different degrees. TCB-2 reduced baseline, anticipatory, and consummatory licking. However, the cue-locked temporal pattern of licking remained visible. TCB-2 therefore appeared to have a greater effect on the amount or vigor of licking than on the temporal organization of the previously learned response. The reduction in licking could reflect changes in motivation, motor output, reward consumption, or physiological state and does not necessarily indicate loss of the learned association. Cue-related pupil dilation also remained evident at the lower doses of TCB-2, and TCB-2 alone produced no statistically detectable changes in eye-position dynamics or blinking. These findings suggest that the mice remained engaged in the task, although some aspects of task performance were clearly affected. Importantly, the present experiments examined the expression of a previously learned association rather than the acquisition of a new association. They therefore cannot determine whether TCB-2 affects learning, extinction, the updating of learned relationships, or cognitive flexibility. Future studies using changes in cue–reward relationships or tasks designed to separate response vigor from associative control will be needed to examine these possibilities.

In freely moving mice, the highest dose of TCB-2 caused a marked reduction in locomotor activity early in the test session, followed by a gradual recovery. Previous studies have found either no significant effect of TCB-2 on open-field activity or an increase in locomotor activity mediated by 5-HT_2A_ receptors (Fox et al., 2010; Halberstadt et al., 2013). Differences in dose, habituation, apparatus, session duration, and the time period used for analysis may explain these inconsistent findings. The time-course analysis in the present study showed that the reduction in locomotor activity was concentrated in the early part of the session. This effect could therefore be overlooked or underestimated when locomotor activity is averaged across the entire session. TCB-2 also reduced fecal and urinary output, showing that its acute effects were not limited to locomotion. These reductions could result from changes in autonomic regulation, stress responses, fluid balance, or the amount of movement. The present experiments cannot distinguish among these explanations. Volinanserin did not clearly prevent the reductions in locomotor activity and fecal or urinary output. These findings therefore provide limited evidence that the observed changes were mediated specifically by 5-HT_2A_ receptors under the conditions tested.

TCB-2 produced a marked, dose-related increase in head-twitch responses, consistent with previous findings (Fox et al., 2010). The head-twitch response is widely used in rodents as a behavioral measure of psychedelic-like drug activity and is strongly associated with 5-HT_2A_ receptor agonism (Halberstadt & Geyer, 2018). The potency of serotonergic hallucinogens in producing this response also correlates with their reported hallucinogenic potency in humans, making the assay useful for comparisons among compounds (Halberstadt et al., 2020). In the present study, head-twitch responses were evident during the first recorded interval and continued throughout the observation period. Volinanserin reduced the response under some conditions but did not consistently abolish it across the TCB-2 doses tested. This result differs from previous studies in which 5-HT_2A_ receptor antagonists strongly suppressed psychedelic-induced head-twitch responses (Halberstadt & Geyer, 2018; Jaster et al., 2022b; Takaba et al., 2024). Several factors could explain the incomplete effect of volinanserin, including its dose and timing, incomplete receptor occupancy, the functional selectivity and broader pharmacological profile of TCB-2, and differences in experimental or analytical methods. The present results clearly demonstrate that TCB-2 induces a robust head-twitch response. However, they provide only limited evidence about the receptor mechanisms underlying this response under the conditions tested.

The different patterns observed across the measured responses show that the head-twitch response alone cannot fully characterize the effects of a serotonergic compound. TCB-2 altered pupil size, licking, locomotor activity, and fecal and urinary output. In contrast, TCB-2 alone produced no statistically detectable changes in eye position or blinking, and the cue-locked temporal pattern of conditioned licking remained evident. Head-twitch counts do not capture these differences and cannot determine whether a compound causes a broad behavioral disruption or has strong effects on only some aspects of behavior and physiology. A recent comparison of lisuride and LSD similarly found that the two compounds produced different patterns of head-twitch, behavioral, physiological, cognitive, and cortical effects (King et al., 2026). Evaluating several measures is therefore important when comparing the pharmacological profiles of serotonergic compounds. Automated detection remains valuable because it provides an objective and scalable method for measuring head-twitch responses. However, its pharmacological value increases when it is used alongside other behavioral and physiological measurements. Continuous monitoring of pupil size, eye position, and blinking is particularly useful because these measures provide information about physiological state and task engagement without interrupting ongoing behavior. The classifier used in the present study also showed good agreement with manual annotations. Nevertheless, testing the classifier using recordings from animals that were not included at any stage of classifier development would provide a more rigorous assessment of its ability to generalize to new data.

The effects of volinanserin differed across the measures examined. Volinanserin partially reduced TCB-2-induced pupil constriction and head-twitch responses under some conditions but did not consistently prevent the changes in licking, locomotor activity, or fecal and urinary output. Several factors could explain this pattern, including differences in the dose–response relationship for each measure, incomplete receptor blockade, pharmacological effects of TCB-2 that were not adequately blocked under the present conditions, and differences between the independently tested cohorts (Fox et al., 2010; Halberstadt et al., 2013; Jaster et al., 2022b). The selectivity and effectiveness of an antagonist depend on its dose, receptor occupancy, timing of administration, and experimental conditions (Casey et al., 2022). The presence or absence of an effect after a single antagonist dose is therefore not sufficient to identify the receptor mechanism responsible for that effect. Differences in age, sample size, and animal-to-animal variability between the cohorts may also have affected the results. It remains unclear whether the observed differences in volinanserin sensitivity reflect distinct receptor mechanisms or differences in pharmacological and experimental conditions. Future studies using multiple doses of volinanserin, direct measurements of receptor occupancy, experimental designs that allow direct comparison of pretreatment conditions, and complementary genetic or circuit-based methods will be needed to distinguish among these possibilities.

The combination of measures used in the present study may also help compare serotonergic compounds with different pharmacological properties. Several compounds have been reported to promote neural plasticity or produce antidepressant-like effects in preclinical models while inducing little or no head-twitch response (Cameron et al., 2021; Cao et al., 2022; Dong et al., 2021). Compounds with hallucinogenic effects and those described as non-hallucinogenic may also differ in their receptor interactions and downstream signaling (Cao et al., 2022; Dong et al., 2021). The present findings do not show that pupil constriction, the stability of eye position, blinking, or any other single measure is a specific marker of hallucinogenic or therapeutic activity. Instead, the overall pattern of behavioral and physiological responses may provide more information than any single measure. Adding pupil dynamics, eye position, and blinking to established measures such as head-twitch responses and locomotor activity may help identify differences between compounds that appear similar when evaluated with only one assay.

Several limitations should be considered when interpreting the present findings. First, only male mice were examined, although sex differences have been reported in head-twitch responses and 5-HT_2A_ receptor signaling (Jaster et al., 2022a). Second, only one dose of volinanserin was tested, and receptor occupancy was not measured. The experiments with and without volinanserin pretreatment also involved independent cohorts that differed in age and sample size. These differences prevent direct comparison between the two experiments and may have contributed to the inconsistent effects of volinanserin across the measures examined. In addition, the absence of statistically significant changes in eye position and blinking following TCB-2 alone does not demonstrate that these measures were unaffected. Small effects may not have been detected because of the sample size and precision of the present analyses. The pharmacology of TCB-2 also limits the conclusions that can be drawn from the antagonist experiments. TCB-2 has high affinity and agonist activity at the 5-HT_2A_ receptor, but different ligands acting at this receptor can activate downstream signaling pathways to different degrees. TCB-2 itself shows functional selectivity across receptor-mediated signaling mechanisms (López-Giménez & González-Maeso, 2018; McLean et al., 2006). Some of its effects may therefore reflect signaling patterns or pharmacological actions that are not shared by all serotonergic psychedelics (Di Giovanni & De Deurwaerdère, 2018; Fox et al., 2010). Under the present conditions, neither the persistence of an effect after volinanserin pretreatment nor its reduction by volinanserin is sufficient to identify the receptor or signaling mechanism responsible. Finally, the present study examined only the acute effects of TCB-2. It therefore cannot determine whether TCB-2 produces delayed or lasting neural and behavioral changes similar to the epigenomic, synaptic, and structural plasticity reported after administration of other serotonergic psychedelics (de la Fuente Revenga et al., 2021; Shao et al., 2021).

In conclusion, acute administration of TCB-2 produced a distinct pattern of behavioral and physiological effects. TCB-2 caused pupillary constriction, reduced licking and locomotor activity, decreased fecal and urinary output, and markedly increased head-twitch responses. However, it did not affect all measures in the same way. The cue-locked temporal pattern of conditioned licking remained evident, and TCB-2 alone produced no statistically detectable changes in eye-position dynamics or blinking. Volinanserin did not consistently prevent the effects of TCB-2, and the present experiments therefore do not establish the receptor mechanism underlying each response. Automated measurement of head-twitch responses, combined with continuous monitoring of pupil dynamics, eye position, blinking, and learned behavior, provides a broader method for comparing serotonergic compounds and detecting pharmacological differences that may not be apparent from a single behavioral assay.

## Ethics Statement

All experimental and animal-housing procedures complied with applicable Japanese national regulations and institutional guidelines for the care and use of laboratory animals. The experimental protocols were reviewed and approved by the Institutional Animal Care and Use Committee of Keio University.

## Author Contributions

MY and KT contributed to the conceptualization and design of the study. MY conducted the head-fixed trace-conditioning experiments with assistance from HI, KH, SM, KY, and KT, and conducted the open-field and head-twitch response experiments. MY performed the behavioral analyses, with IA contributing to the manual validation of head-twitch responses and JM contributing to the automated head-twitch analysis. MY and KT prepared the original draft and created the figures. MY, HI, KH, SM, KY, IA, JM, and KT contributed to interpretation of the findings and critical revision of the manuscript. KT supervised the study. All authors reviewed and approved the final manuscript.

## Acknowledgements

The authors thank Haruki Kasahara, Shunsuke Nakajima, and Shudo Yoshida for their assistance with animal care. This work was supported by JSPS KAKENHI grant numbers 23H02787, 23K27478, 23K22376, and 24H00729 to KT; 24K16869 and 24KJ0069 to KY; and 24K06626 and 25KJ0306 to KH. Additional support was provided by the Keio Academic Development Fund, the Keio Gijuku Fukuzawa Memorial Fund, the SR Foundation, and the HOKUTO Foundation for the Promotion of Biological Science to KT. During preparation of this manuscript, the authors used ChatGPT (OpenAI) solely to improve the clarity and readability of the text. The tool was not used to generate, analyze, or interpret the experimental data. All AI-assisted text was reviewed and edited by the authors, who take full responsibility for the accuracy and integrity of the final manuscript.

## Data Availability Statement

The data supporting the findings of this study and the custom code used for the analyses are available from the corresponding author upon reasonable request.

